# Conserved spatiotemporal expression landscape of dominant tRNA genes in human and mouse

**DOI:** 10.1101/2023.08.14.553175

**Authors:** Evan Y. Wu, Laura Landry

**Affiliations:** Memphis University School, 6191 Park Ave, Memphis, TN 38119, USA

## Abstract

Transfer RNAs are integral for protein synthesis and the interpretation of the information contained in DNA. To date, a few methods, including custom microarrays and custom targeted sequencing, have been used to quantify tRNA. However, methods using available RNA-sequencing data have not yet been reported. We created a bioinformatics pipeline to quantify the highly expressed tRNAs in RNA-Seq effectively, demonstrated by the preserved ratio of the expression levels of two massively duplicated tRNA^Ala^ genes in mouse. Using this quantification, we examined the tRNA expression with relation to tissue type and developmental stage in both human and mouse. Heart exhibited the highest overall tRNA expression for both human and mouse. Furthermore, tRNA expression grew to a peak before decreasing steadily with developmental stage, a trend that was conserved in both human and mouse. The two mitochondrial tRNA genes, tRNA^Ser^(TCA)(m) and tRNA^Leu^(TTA)(m), which partly contribute to these trends, have been attributed to various human diseases. The tissue-specific high expression of tRNA^Gln^(CAG) and tRNA^Gln^(CAA) in human brains, especially in hindbrain and cerebellum, suggests their important roles in neurological disorders. In summary, our approach revealed conserved spatiotemporal expression of highly expressed tRNAs in both human and mouse. Our method can be applied to other RNA-Seq data to examine the roles of these tRNAs in different human diseases or scientific studies.

## Introduction

Transfer RNAs, or tRNAs, exist in all organisms and serve as the universal interpreter of the genetic information of DNA. There are qualitative and quantitative variations for various tRNA genes. Qualitatively, numerous copies of the same tRNA genes may be duplicated on the genomes of different species, with different copies accumulating mutations during evolution. Quantitatively, not all copies of tRNA genes are always expressed or present in every tissue. Mutations and defects in tRNAs may cause human diseases [1-3]. For example, several tRNA mutations are associated with hypertension [4]. The mitochondrial tRNA^Leu^(TTR) mutation may lead to MELAS (mitochondrial encephalomyopathy, lactic acidosis, and stroke-like episodes), a disease attacking mainly the nervous system and muscles in young adults [5]. Similarly, another mutation on the mitochondrial tRNA^Lys^ gene causes MERFF (myoclonic epilepsy with ragged-red fibers), a multisystem disease that also primarily affects children and young adults [6], [7]. Likewise, aberrant tRNA expression is attributed to a wide variety of human diseases. For example, overexpression of tRNA is connected with breast cancer [1], [3]. On the other hand, downregulation of the tRNA^Ser^(TCY) is related to SARS2- related diseases [8]. Therefore, characterization of the heterogeneity and dynamics of tRNA expression and variations is important to understand how the cellular translational machinery contributes to human diseases. In this study, we plan to focus on the dynamics of tRNA expression levels in different tissues and developmental stages.

Thus far, a few tRNA databases have documented the number of tRNA copies in the human (tRNAdb: 359, GtRNAdb: 429) and mouse (tRNAdb: 14, GtRNAdb: 400) genomes [9, 10]. In addition, RepeatMasker has annotated all homologous tRNA genes [11], including pseudogenes, under RNA repeats, regardless of their function and expression. There have been multiple methods for quantifying tRNA. During the 1980s, researchers relied on Northern blot to measure tRNAs. In recent years, high-throughput technologies started playing a crucial role in characterizing the tRNA pools. For instance, microarrays with 63 probes (42 for nuclear tRNAs and 21 for mitochondrial tRNAs) were used to compare tRNA levels for humans [12], but one limitation of this method is the difficulty of determining the precise locus of contributing tRNA genes on the genome. With microarray, Dittmar *et al* has shown that tRNA levels and codon usage vary across different tissues [12]. However, it lacks the granularity of tRNA tissue specific expression due to the limitation of the assay. Furthermore, the variance in tRNA levels and codon usage of different tissues was only considered in humans. RNA-sequencing is a much better option for analyzing tRNA because it not only determines the expressed tRNA loci, but it also determines variations in the tRNAs compared to the reference genome. However, most tRNAs do not have poly-A tails, so RNA-Seq based on poly-A enriched protocol cannot be used to measure tRNA expression [13]. To date, tRNA levels have not been investigated in relation to developmental stages.

Aside from nuclear tRNA genes, mitochondrial tRNA levels varied significantly across 5 different mouse tissues (brain, heart, liver, skeletal muscle, and kidney) with heart having the highest tRNA levels and liver having the lowest [14]. However, mitochondrial tRNA differences across tissues in humans have not been thoroughly explored.

In this study, we leveraged previously generated total-stranded RNA-seq data for two mammalian species in various tissues and different developmental stages [15]. We characterized the spatiotemporal landscape of dominant tRNA genes to understand the trends and patterns of expressed tRNAs on different human and mouse tissues along with commonality and differences between human and mouse. Specifically, we ask the following questions: 1. Does overall tRNA abundance vary across tissues? If so, is it the same pattern in human and mouse? 2. Does overall tRNA abundance vary by developmental stages? Is it the same pattern in human and mouse? 3. Does individual tRNA express at the same level? If not, is it the pattern the same in human and mouse?

## Results

### Quantification of tRNA gene expression and normalization for fair comparison

To characterize the dynamics of tRNA accurately in different tissues and developmental stages, we analyzed the RNA-seq dataset that was comprehensively performed for hundreds of people and mice in a recent study [15]. We described the workflow in **S1 Fig**.

The mouse samples were collected from e10.5 to 9wpb, and the human samples were collected from fetus to adult (**S1 Table**). For all samples except human ovary, we have more than 20 samples (**S2A Fig**). For mouse, we had 160 male samples and 157 female samples, so the sex is well-balanced. However, for human, we had 203 male samples and 109 female samples, demonstrate some bias toward males. In total, we had 317 samples for mouse and 312 samples for human. We included forebrain and hindbrain for human in early development.

The total stranded RNA-seq protocol in this study allows us to identify tRNA genes with and without poly-A tails. Briefly, we mapped the sequencing read data across brain, cerebellum, heart, kidney, liver, ovary, and testis tissues (**S1 Fig**) and counted the total number of sequencing reads to use as the denominator for normalization (**S1 Table**). We extracted mapped sequencing reads from all genomic loci that were annotated as tRNAs by RepeatMasker [11]. We used RepeatMasker instead of other databases since the tRNA annotated by RepeatMasker is the most comprehensive (**S2 Table**). We noted that the extracted reads from these loci were not always perfectly matched to the tRNA reference sequences. To avoid overcounting, we extracted the reference tRNA sequences, performed a BLAST search against the unique sequencing reads, and only retained the reads with at least 50bp perfectly overlapping with reference tRNA sequences without gaps as the estimate of the expression level. This dramatically reduced the counts of reads (**S3 Table**).

One challenge to comparing the tRNA abundance across samples is that different sequencing depths lead to different numbers of total reads (**S2B Fig**). To combat this, we normalized both sets of data with the total number of reads instead of the total number of reads mapped to tRNA genes (**S2C Fig**). This seems to be an effective approach as the variance in total tRNA abundance within each tissue type significantly reduced after normalization (data not shown).

### Small fraction of tRNA genes is detected as highly expressed by RNA-Seq

We defined a tRNA as “highly expressed” if it has at least three samples with > 1 TPM (<u>T</u>ransfrags <u>P</u>er <u>M</u>illion reads) after normalization with total reads per sample. We only detected 12 mouse tRNA genes and 19 human tRNA genes meeting this criterion after merging the counts from various loci of tRNA gene copies. Notably, the top two high-expression tRNA genes are mitochondrial tRNA^Ser^(TCA) and tRNA^Leu^(TTA) in both human and mouse. Also, tRNA^Ala^(GCA) and tRNA^Ala^(GCY) are only highly-expressed in mouse but not in human, which is very likely due to the excessive duplication of Ala-tRNA genes in the mouse genome (3,536 copies, **S2A Table**). Among them, there are 749 copies of tRNA^Ala^(GCA) and 2,588 copies of tRNA^Ala^(GCY) on mouse genome, which matches well with the overall expression level that we observed for these two tRNA genes (average count ratio: tRNA^Ala^ GCY/GCA=3.46±0.77, **S3A Table**).

We suspect that most tRNA genes were undetectable using the current RNA-seq protocol, likely due to the size selection in the RNA-seq library preparation, which focused on relatively long RNA molecules (> ∼ 100bp), while most tRNA genes are about 76 bp. Therefore, this technology could be biased to tRNA genes with relatively long unprocessed precursors or mitochondrial tRNAs. Nevertheless, the concordance of the tRNA gene copies (tRNA^Ala^ GCY/GCA=3.45) and detected tRNA read counts for mouse tRNA^Ala^ is convincing. Thus, we can examine the general trends in these highly expressed tRNA genes.

### Total tRNA pools vary by tissues and developmental stages in mouse and human

First, we investigated the total tRNA abundance in human and mouse with relation to tissue types. For mouse, heart tissue exhibited the highest overall tRNA abundance (**Fig 1A**). However, for human, the total tRNA abundance was high not only in heart, but also in brain and kidney tissues (**Fig 1B**). Notably, the tRNA gene expression in heart and ovary in mouse and human showed bimodal distributions, suggesting additional sources of variability. Next, we broke down the changes in the total tRNA abundance across tissues over different developmental stages for both mouse and human. For mouse, tRNA abundance in heart tissue increased steadily over time (**Fig 1C**). Kidney tissue total tRNA abundance skyrocketed at around 2wpb, then dropped sharply (**S3D Fig**). For human, heart, liver, brain, and kidney, tRNA abundance increased sharply from infant to child (**Fig 1D**). The levels peaked during childhood, before dropping slightly at the teenage stage (**Fig 2D**). On the other hand, cerebellum tRNA abundance remained relatively constant. Since data for humans comes from deceased humans, our information is limited for some tissues in children, teens, and adults. Nevertheless, the variance of the total tRNA abundance estimates for the same condition is relatively small, confirming that the general trends likely hold true.

**Figure 1.**
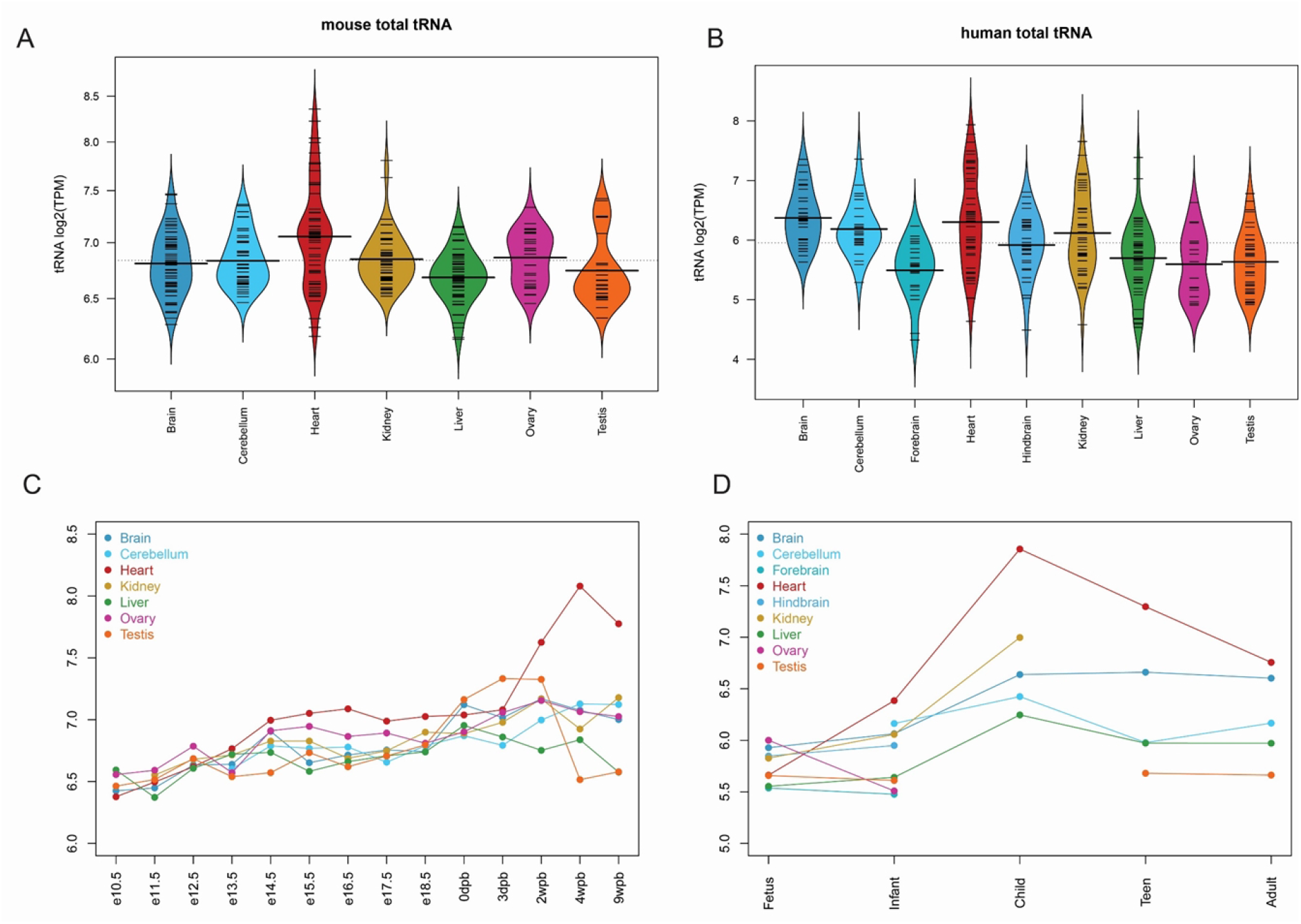
The total tRNA abundance varies across tissues in both mouse and human. **A**) The total tRNA abundance is higher in heart in mouse after normalization to the whole transcriptome; **B**) The total tRNA abundance is higher not only in heart but also in whole brain (except forebrain), liver, and kidney in human after normalization to the whole transcriptome. **C**) The total tRNA abundance shows different dynamics between heart and other organs during the mouse development. **D**) Heart total tRNA expression also peaked at early childhood in human. Increase in total RNA expression in early childhood is also observed in brain, kidney, and liver in human.

**Figure 2.**
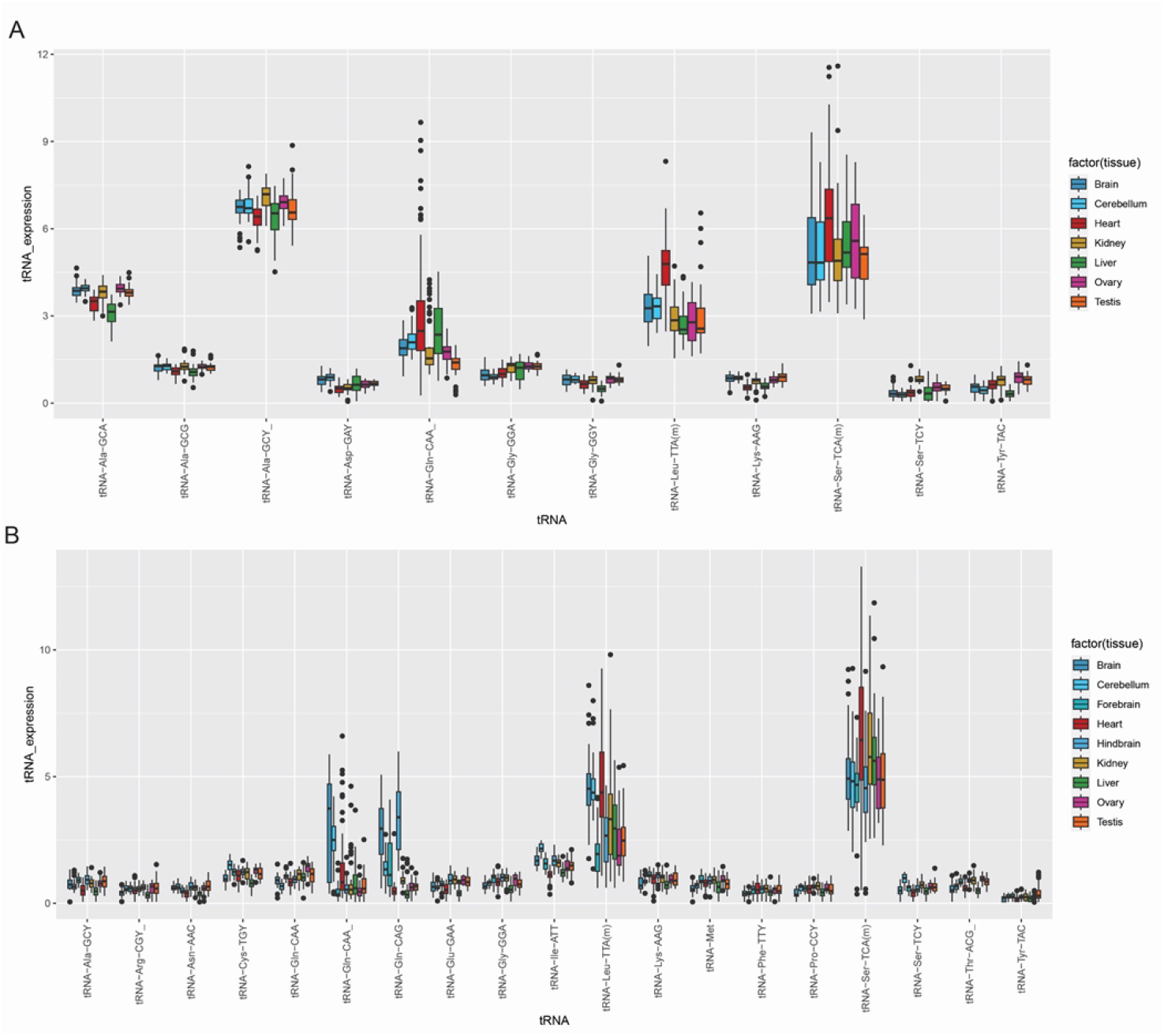
The expression of individual mouse tRNA varies across tissues and developmental stages. **A)** The tRNA expression for different amino acids varies by tissues, with heart expression being notably high for tRNA^Gln^(CAA), tRNA^Leu^(TTA)(m), and tRNA^Ser^(TCA)(m), and liver expression being high for tRNA^Gln^(CAA). **B)** The tRNA expression for different amino acids varies by human tissues, with heart expression being notably high in tRNA^Leu^(TTA)(m), and tRNA^Ser^(TCA)(m). The high expression in brain, especially hindbrain and cerebellum, is obvious for tRNA^Gln^(CAA), tRNA^Gln^(CAG) and tRNA^Leu^(TTA)(m).

### Individual tRNAs for some amino acids show tissue specific expression pattern

For mouse, the tRNA gene with the highest expression is tRNA^Ala^(GCY), followed by tRNA^Ser^(TCA)(m), tRNA^Ala^(GCA), tRNA^Leu^(TTA)(m), and tRNA^Gln^(CAA) (**Fig 2A**). The high expression of mouse tRNA^Ala^ did not exhibit many tissue-specific patterns, except slightly lower expression of tRNA^Ala^(GCA) in liver and higher expression of tRNA^Ala^(GCY) in kidney. The high tRNA gene expression in heart seems to be driven by tRNA^Gln^(CAA), tRNA^Leu^(TTA)(m), and tRNA^Ser^(TCA)(m) (**Fig 2A**), two of which are mitochondrial tRNAs. Interestingly, although overall liver tRNA expression tends to be generally low in mouse (**Fig 1A**), tRNA^Gln^(CAA) is much more highly expressed in liver than in other tissues (**Fig 2A**). Additionally, mouse tRNA^Ser^(TCY) is expressed at relatively higher level in kidney when compared with other tissues. In general, these expressed mouse tRNAs are highly expressed in brain and cerebellum.

Similarly, among the 19 human tRNA genes that are detectable, the most highly expressed tRNA is tRNA^Ser^(TCA)(m), followed by tRNA^Leu^(TTA)(m), tRNA^Gln^(CAG) and tRNA^Gln^(CAA) (**Fig 2B**). Unlike in mouse, human tRNA^Ala^ is not highly expressed. As before, the two mitochondrial tRNAs, tRNA^Leu^(TTA)(m), and tRNA^Ser^(TCA)(m), are the major contributors to high heart tRNA abundance (**Fig 2B**). In contrast with mouse, there is much higher expression of tRNA^Gln^(CAA), tRNA^Gln^(CAG), and tRNA^Leu^(TTA)(m) in human whole brain and cerebellum (**Fig 2B**).

### The expression of certain individual tRNAs fluctuated during organ development

In mouse, most of the 12 detectable tRNA genes, including the highly expressed tRNA^Ala^(GCA), are expressed with little change at different developmental stages (**Fig 3A**). However, tRNA^Ser^(TCA)(m), tRNA^Leu^(TTA)(m), and tRNA^Gln^(CAA) showed increased expression with organ development, especially after birth (**Fig 3A**). In contrast, tRNA^Ala^(GCY) showed slightly lower expression in early mouse embryos and after birth (**Fig 3A**).

**Figure 3.**
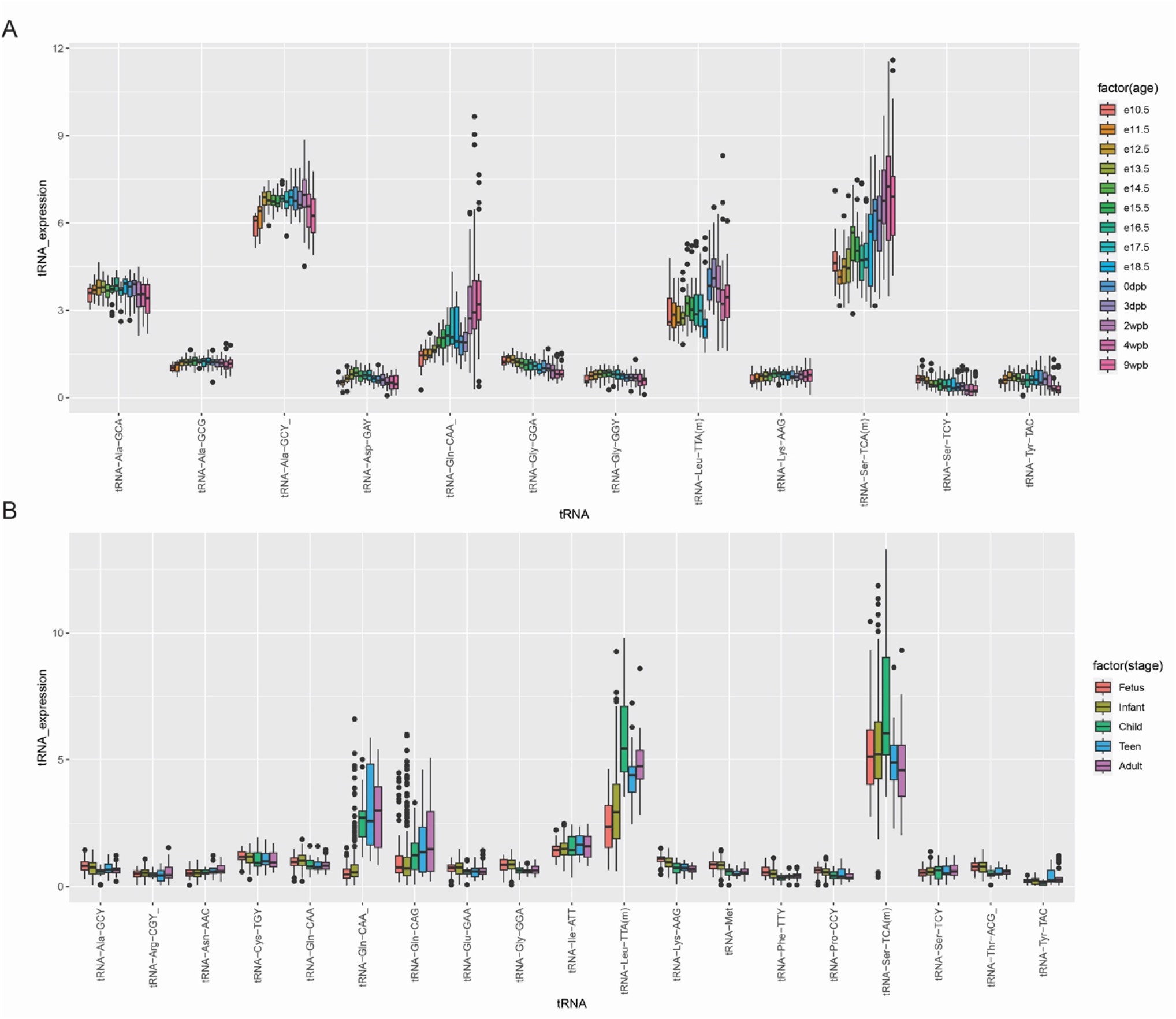
The expression of individual tRNA varies by developmental stages. **A**) mouse tRNA expression for certain amino acids varies over developmental stages. For tRNA^Gln^(CAA), tRNA^Leu^(TTA)(m), and tRNA^Ser^(TCA)(m), there is a general increase over time. tRNA^Ala^(GCY) shows an increase in expression followed by a decrease. **B**) human tRNA expression for certain amino acids varies over developmental stages. tRNA^Leu^(TTA)(m), and tRNA^Ser^(TCA)(m) peaked at childhood. there is a general. tRNA^Gln^(CAA) and tRNA^Gln^(CAG) showed higher expression in childhood and adulthood.

For the 19 detectable human tRNA genes, the expression of two mitochondrial tRNAs, tRNA^Ser^(TCA)(m) and tRNA^Leu^(TTA)(m), which are expressed at the highest level in heart (**Fig 2B**), appears to peak at early childhood development (**Fig 3B**). The two brain-specific tRNA^Gln^(CAA) and tRNA^Gln^(CAG) showed much higher expression in childhood and adulthood when compared with fetus and infant (**Fig 3B**). Similarly, higher expression of tRNA^Gln^(CAA) was also observed in mouse after birth (**Fig 3A**), although this tRNA does not seem to be expressed specifically highly in mouse brains as is the case in humans (**Fig 2A**).

## Discussion

Transfer RNA plays a central role in the biological processes of all cells. However, the understanding of its abundance and dynamics is surprisingly limited. Although there were a few early efforts to quantify the expression of tRNAs using high-throughput technologies, it remains unclear if we can extract any useful information about tRNAs from the large amount of publicly available RNA-Seq data. In this study, we made the first attempt to explore the feasibility of quantifying highly expressed tRNAs in a wide variety of human and mouse tissues at different developmental stages. In our computational pipeline, we first employed standard STAR mapping against both the human and mouse genome with Genecode annotation to exclude the reads that map to protein-coding genes and other long non-coding RNA genes. Then we specifically extracted reads mapped to all computationally-predicted tRNA genes from RepeatMasker. We used this exhaustive approach to avoid missing any regions that are related to tRNAs. To increase the accuracy of the quantification, we applied a pairwise alignment approach to ensure that only reads matching perfectly to reference tRNA sequences (no mismatches, no gaps) were retained, as multiple tRNA gene copies could differ by only a few bases. Although we might miss mutant tRNAs in some of these samples, the major trends of the tRNA dynamics should remain true since we relied on over 300 samples in each species. A few lines of evidence support the validity of this approach. First, we only observed high tRNA^Ala^ expression in all mouse samples but not in any human samples. This is consistent with the fact that tRNA^Ala^ gene duplication only occurs in the mouse genome but not in the human genome. Second, the expression level of two dominant tRNA^Ala^ in all the mouse samples preserves the ratio of the gene copies of the two tRNA^Ala^, indicating that the expression level is proportional to gene copies. Third, although mitochondrial genes are the most highly expressed tRNA genes, their expression levels are still lower than the expression of tRNA^Ala^(GCY), which is consistent with the fact that there are only about 300 to 400 copies of mitochondrial copies per cell. Finally, the conserved patterns found for these genes across species demonstrate that this approach captures the actual biological signals in these cells.

In both human and mouse, we found overall tRNA abundance to be highest in heart. This is an unreported and novel discovery. We also verified that, in general, the tRNA expressions in liver, testis, and ovary are lower than that of brain tissue, which is consistent with previous reports [12]. Furthermore, our findings that the expression levels of mitochondrial tRNA^Leu^ and tRNA^Ser^ in ovary are less than in brain match with previous findings [12]. For mouse, the kidney tRNA expression was highest in tRNA^Ala^(GCY). However, since human tRNA expression is low for tRNA^Ala^, this pattern is not observed in human. Clearly, our approach relies on the absolute expression level of each tRNA, so it is easier to interpret the results.

We also discovered that tRNA expression level changes with developmental stage in both human and mouse. For most tissues, the tRNA expression increases, before peaking soon after birth for both human and mouse. An exception is kidney tissue, which demonstrates a consistent increase with development for both human and mouse. Overall, the temporal relationship between tRNA expression and developmental stage is consistent between human and mouse. One limitation of our observation is that we don’t have the full representation of human tissues at exactly the same developmental stages as biological replicates, so the exact time point of tRNA gene expression changes is difficult to determine.

The understanding of tissue-specific and temporal expression patterns of tRNAs is important to comprehend the etiology of human diseases. Mutations on the mitochondrial tRNA^Leu^(TTR) have been observed to cause diseases, including MELAS and hypertension [4, 5]. This is consistent with the high expression of the housekeeping tRNA^Leu^(TTR) gene in brain and heart tissues in both human and mouse. Additionally, changes in the mitochondrial function of the tRNA^Ser^(TCN) have been linked to a DNA mutation associated with deafness and SARS2-related diseases [8, 16]. The housekeeping tRNA^Ser^ (TCA)(m) exhibits high levels in liver, heart, and brain tissues, confirming its importance in multiple organs. Notably, we notice that tRNA^Gln^ is highly expressed in human brains, in particular, hindbrain in early development and cerebellum in adult. Therefore, we hypothesized that the mutation in tRNA^Gln^ should also lead to human diseases. Indeed, we found two reports connecting the tRNA^Gln^ mutation A4336G to Parkinson disease or migraine [17, 18]. In summary, our approach appears to be robust in quantifying high-expression tRNAs from total stranded RNA-Seq. Although we have not tested its validity for poly-A enriched library preparation-based RNA-Seq, we believe that the catalog of these high-expression tRNAs across normal and diseased tissues would greatly help to understand the metabolism-related molecular mechanism of human diseases.

## Materials and Methods

### Data download

The raw sequencing data was downloaded from EGA ArrayExpress (https://www.ebi.ac.uk/biostudies/arrayexpress). For the whole transcriptome sequencing (RNA-Seq) data generated in the study of Cardoso-Moreira et al (2019) [15], only human (E-MTAB-6814) and mouse (E-MTAB-6798) datasets were used in this study (**S1 Table**). Due to the limited availability of human samples, we grouped multiple samples into five main categories: fetus, infant, child, teen, and adult. Although we might lose the resolution, the major trend observed should hold true with. All RNA-Seq were done with total stranded library preparation protocol, which captures tRNAs without poly-A tails.

For tRNA annotation, we examined three databases: tRNAdb [9], GtRNAdb [10] and RepeatMasker [11]. The repeatmaker track was downloaded from UCSC genome browser [19]. Since whole transcriptome sequencing can unbiasedly characterize all transcripts regardless of annotation, we decided to investigate the largest of collection of tRNA annotation in RepeatMasker for completeness (**S2 Table**). But for the comparison, we focused on the expressed tRNA genes only.

### Mapping of whole transcriptome sequencing data and quantification of gene expression

We mapped the RNA-Seq data against reference genomes (human: GRCh38, aka, hg38; mouse: GRCm38, mm10) with STAR aligner (2.7.9a, [20]). The following parameters were used in the mapping: “--limitBAMsortRAM 67108864000 –genomeDir STAR-index/2.7.9a --outFilterMultimapNmax 20 --alignSJoverhangMin 8 -- alignSJstitchMismatchNmax 5 -1 5 5 --alignSJDBoverhangMin 10 -- outFilterMismatchNmax 999 --outFilterMismatchNoverReadLmax 0.04 --alignIntronMin 20 --alignIntronMax 100000 --alignMatesGapMax 100000 --genomeLoad --outSAMmapqUnique 60 --outSAMmultNmax 1 --outSAMstrandField intronMotif -- outSAMattributes NH HI AS nM NM MD --outSAMunmapped Within --outSAMtype BAM SortedByCoordinate --outReadsUnmapped None --outSAMattrRGline -- chimSegmentMin 12 --chimJunctionOverhangMin 12 --chimSegmentReadGapMax 3 -- chimMultimapNmax 10 --chimMultimapScoreRange 10 --chimNonchimScoreDropMin 10 --chimOutJunctionFormat 1 --quantMode TranscriptomeSAM GeneCounts -- twopassMode Basic --peOverlapNbasesMin 12 --peOverlapMMp 0.1 --outWigType wiggle --outWigStrand Stranded --outWigNorm RPM”. Gencode annotation was used for gene model in the mapping for splicing events (human: r31; mouse: M22. [21]).

The total number of mapped reads was obtained with samtools [22]. The mapping metrics were summarized in **S1 Table**. The reads mapped to annotated tRNA loci were extracted with samtools. To retain the high accuracy counts, all the extracted reads were merged as unique sequences and serves as the database (per sample) for BLAST search. Each of the annotated reference tRNA sequence was BLAST against these databases. Only hits with no gap, and more than 50bp of perfect matches were retained to summarize the counts. Finally, the counts for each tRNA gene with the same anticodon were aggregated to allow comparison (**S3 Table**). The aggregated counts were normalized by the total number of sequencing reads per sample for fair comparison across samples.

### Statistical analysis and visualization

All statistical analyses and visualization were performed under R environment (R/4.1.0, https://www.r-project.org/). Various R packages (“ggplot2”, “beanplot”) were used to help visualization using bean plots, boxplots, or heatmap.

## Supplementary Information

### Supplementary Figures

**S1 Figure. The workflow of data processing and analysis described in this study. A)** The raw Illumina RNA-Seq data as mated pairs (E-MTAB-6798 and E-MTAB-6814) were downloaded from the EGA ArrayExpress database. The stranded RNA-Seq data was mapped against the human genome (GRCh38, or hg38) and the mouse genome (MCG38 or mm10) with STAR mapping, using Gencode annotation for gene coding regions. The total mapped reads were extracted with samtools to be used as the denominator for normalization. The mapped reads at tRNA gene loci annotated in RepeatMasker program were extracted with samtools. The reads were further merged into unique sequences with counts, with only sequences with more than 50bp perfectly matching tRNA sequences without gaps kept for the downstream analyses. Multiple reference tRNAs sharing the identical sequences were counted only once to avoid overcounting. Then the reads matched to all tRNA genes coding for the same anticodon were combined as the estimate of expression level. After normalizing with total sequencing reads, the transcript per million reads (TPM) were used in the downstream comparison. The expressed tRNAs (> 1 TPM in at least three samples) were used for comparison of tissue or age-specific expression to ensure meaningful interpretation. **B**) Example merged unique sequencing reads for tRNA^Ser^(TCA)(m) in a human heart sample. The first row is the human reference sequence with the portion annotated as tRNA highlighted in blue. Three unique read sequences with the number of counts (x180, x51, x97) were shown, with extra variable non-template poly-A extensions, highlighted in red, not matching reference sequences. **C**) Three example sequencing reads out of 180 reads with unique read ID from sample “1437sTS_Human_Heart_11w_Male”, contributing to seq_8647. **D**) Highly correlated counts of two major tRNA^Ala^ in mouse samples, preserving the ratio of the number of gene copies on mouse genome.

**S2 Figure. Data quality control and normalization for fair comparison for mouse (left panels) and human (right panels). A)** Number of samples per tissue used analyzed in this study. Most have more than 20 samples, except human ovary, allowing tissue level summary of general trends. **B**). The total number of sequencing reads are highly variable across tissues and samples. Therefore, normalization of tRNA gene expression with total number of reads is necessary to ensure fair comparison across samples and tissues. **C**) After normalizing against total sequencing reads in each samples and gene length (TPM), most tissues have small ranges of total tRNA expression level. Heart tends to be more variable for biological reasons (see **Fig 1**, bimodal distributions)

**S3 Figure Selected tissue specific dynamics of total tRNA abundance in mouse and human.** The temporal increase in overall tRNA abundance by developmental stages in mouse heart (**A**), brain (**B**), liver (**C**), and kidney (**D**). For mouse and human, there is a general increase for all tissues. However, for all tissues except for kidney, there is a jump in expression at birth, with a decrease in expression afterwards.

### Supplementary Tables

**S1 Table The mapping metrics of each RNA-Seq sample used in this study and the sample meta-data**

**S2 Table The annotation of each tRNA from RepeatMasker in reference genomes. A)** mouse genome GRCm38 (mm10). **B**) human GRCh38 (hg38).

**S3 Table The raw counts obtained for each tRNA shared the same anticodon per sample**. **A**) mouse genome GRCm38 (mm10). **B**) human GRCh38 (hg38).

## Notes

### Competing Interest Statement

The authors have declared no competing interest.

